# Small Intestinal Resident Eosinophils Maintain Gut Homeostasis Following Microbial Colonisation

**DOI:** 10.1101/2021.01.30.428930

**Authors:** K. Shah, A. Ignacio, J. Bernier-Latmani, Y. Köller, G. Coakley, M. Moyat, R. Hamelin, F. Armand, N. C. Wong, H. Ramay, C.A. Thomson, R. Burkhard, A. Dufour, T.V. Petrova, N.L. Harris, K. D. McCoy

## Abstract

Intestinal homeostasis following postnatal microbial colonization requires the coordination of multiple processes including the activation of immune cells, cell-cell communication, the controlled deposition of extracellular matrix, and epithelial cell turnover and differentiation. The intestine harbors the largest frequency of resident eosinophils of all homeostatic organs, yet the functional significance of eosinophil recruitment to this organ has long remained enigmatic. Eosinophils are equipped to both respond to, and modify, their local tissue environment and thus are able to regulate the adaption of tissues to environmental changes. We report a critical role for eosinophils in regulating villous structure, barrier integrity and motility in the small intestine. Notably, the microbiota was identified as a key driver of small intestinal eosinophil activation and function. Collectively our findings demonstrate a critical role for eosinophils in facilitating mutualistic interactions between host and microbiota and provide a rationale for the functional significance of their early life recruitment in the small intestine.

**HIGHLIGHTS:**

i. The microbiota is a critical regulator of eosinophil activation and turnover
ii. Eosinophils uphold intestinal barrier integrity following microbial colonization
iii. Loss of eosinophils at steady-state results in increased villous blunting and altered intestinal motility

## INTRODUCTION

Eosinophils are multi-functional granulocytes commonly identified as effector cells mobilised during type II immune responses. Early studies on eosinophils have largely focused on their response to helminth infection or contribution to pathophysiological processes during allergic disorders, including asthma and eosinophilic esophagitis (Huang and Appleton, 2016; Rothenberg and Hogan, 2006). In contrast, the relevance of eosinophil residence in tissues under homeostatic conditions remains elusive (Shah et al., 2020; Weller and Spencer, 2017). Accumulating evidence supports a role for eosinophils in promoting organogenesis/morphogenesis in the developing mammary gland and lung (Gouon-Evans et al., 2000; Loffredo et al., 2020), aiding the repair of injured muscle (Heredia et al., 2013) and liver (Goh et al., 2013), shedding of the uterine lining (Vicetti Miguel et al., 2017) and contributing to metabolic homeostasis in lean adipose (Qiu et al., 2014; Wu et al., 2011).

Out of all the homeostatic organs where tissue-resident eosinophils are found, the gastrointestinal tract harbours the largest population. They are recruited during late gestation and in early life in a manner independent of the microbiota (Jimenez-Saiz et al., 2020; Mishra et al., 1999). Recent studies have provided evidence for an important role of resident eosinophils in immunoregulation and maintenance of immune populations in the gastrointestinal tract, particularly in the small intestine (SI), both during homeostasis (Chu et al., 2014b; Jung et al., 2015; Sugawara et al., 2016) and upon bacterial infection (Arnold et al., 2018; Buonomo et al., 2016). Evidence that tissue-resident eosinophils in the SI exhibit specific characteristics, such as prolonged survival (Carlens et al., 2009), altered-surface marker expression (Chu et al., 2014a), and signs of degranulation (Kato et al., 1998), suggests that these cells are regulated by the local tissue microenvironment. Yet the factors that regulate their residency or activation within the SI have not been defined, nor is it known whether eosinophils impact on the normal physiological function of the SI.

We report an important functional consequence of eosinophil residency in the SI and demonstrate that eosinophils play a critical role in maintaining tissue homeostasis by regulating the extent of microbiota-induced alterations in the SI. We identify the microbiota as a key regulator of SI eosinophil activation, heterogeneity and function and show that eosinophil-deficiency leads to maladaptive changes in small intestinal architecture and physiology upon microbiota colonization, characterized by significant villous blunting, barrier leakage and altered peristalsis.

## RESULTS

### Distribution and localization of resident eosinophils in the small intestine under homeostatic conditions

In order to understand the contributions of eosinophil residency in the SI during homeostasis, we characterized their distribution along its length by flow cytometry. In accordance with previous studies, eosinophils could be identified as SSC^high^CD45^+^Siglec-F^+^ cells (Carlens et al., 2009; Chu et al., 2014a) with high surface expression of CD11b and intermediate levels of CCR3, CD44, CD103, CD11c, Ly6C and Gr1 (Figure 1A). Corroborating previous reports (Chu et al., 2014a), eosinophils were highly enriched in the duodenum and jejunum in comparison to the ileum (Figure 1B). To gain insight into the functions of resident eosinophils, we analyzed their localization within the SI villi using a whole-mount immunostaining protocol (Bernier-Latmani and Petrova, 2016). Eosinophils were found throughout the SI lamina propria and were found in close proximity to stromal cells, such as α-smooth actin (α-SMA)^+^ myofibroblasts (Figure 1C, Figure S1A), enteric glia (Figure 1C) and neural processes, including neuronal axons (Figure S1B). Eosinophils were distributed homogenosly throughout the villus and crypt zones, however were absent in the muscularis mucosa under homeostatic conditions (Figure 1D, Figure S1C).

**Figure 1.**
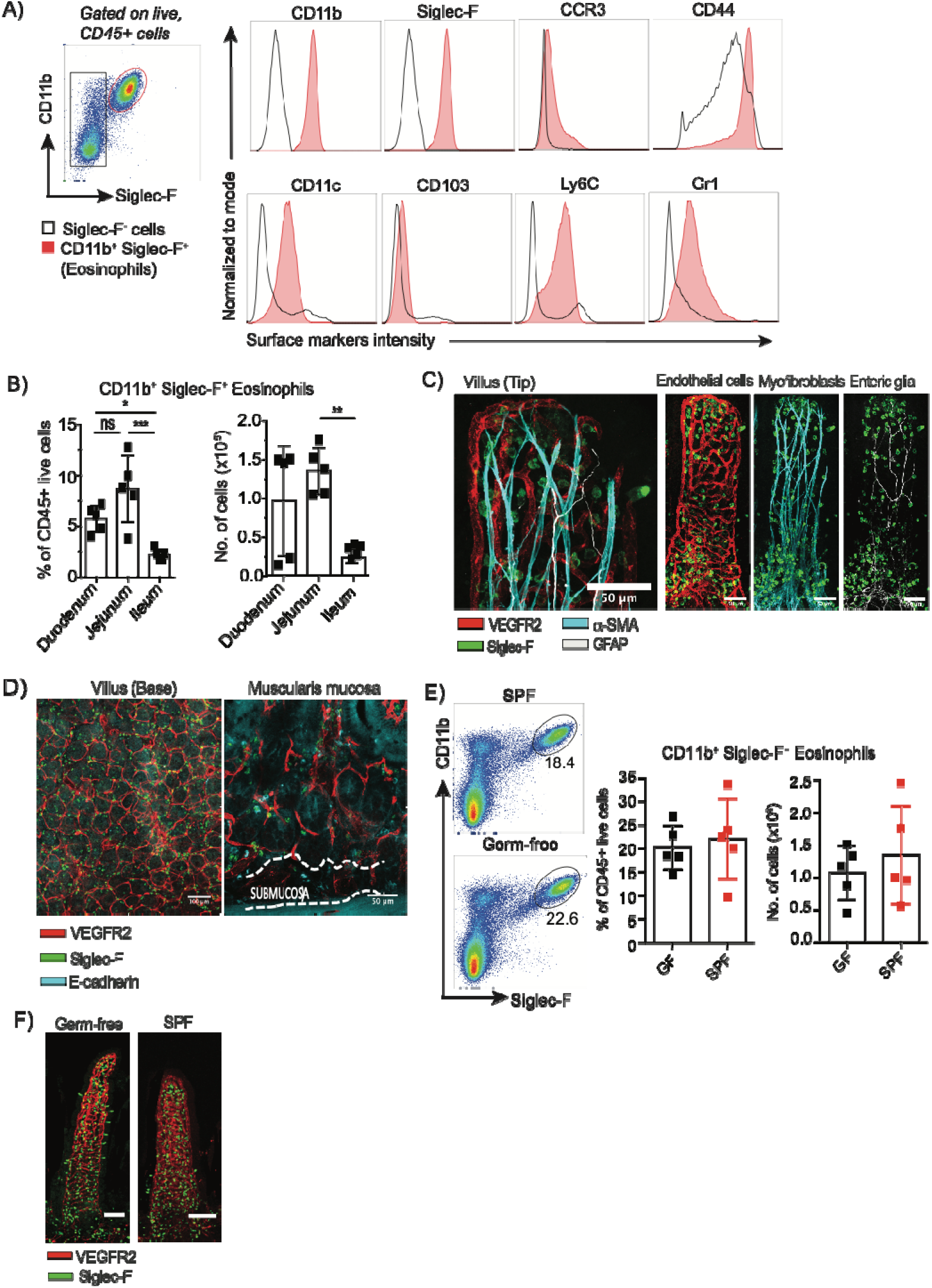
SI eosinophils are localized primarily in the villous lamina propria and are recruited in a microbiota-independent manner. (A, B) Flow cytometry analysis of eosinophils isolated from the SI lamina propria showing (A) Gating strategy, mean fluorescence intensity (MFI) of surface markers, and (B) frequency of eosinophils within total CD45^+^ cells plus absolute eosinophil numbers within distinct SI regions. (C-D) Whole-mount staining of jejunum from BALB/c mice showing (C, from left to right) villous tip view with staining for eosinophils (Siglec-F, green), blood endothelial cells (VEGFR2, red), myofibroblasts (α-SMA, cyan) and enteric glia (GFAP, white); (D, from left to right) muscularis mucosa and crypt view. (E) Flow cytometry analysis of eosinophils isolated from the SI lamina propria of germ-free (GF) or specific-pathogen free (SPF) BALB/c mice showing frequency of eosinophils within total CD45^+^ cells and absolute eosinophil numbers. (F) Whole-mount staining of jejunum from GF or SPF BALB/c mice showing villous tip view with staining for eosinophils (Siglec-F, green) and blood endothelial cells (VEGFR2, red). Flow cytometry experiments show one experiment and are representative of four independent experiments performed using GF and SPF mice from two different animal facilities. For all panels statistical analysis was performed using a student t-test; p<0.05*, p<0.01**, p<0.001***, N.S. non-significant, each bar graph representing data mean ± SD of n=3-5 mice per experiment with individual data points representing a single mouse. See also Figure S1.

Next, we analyzed eosinophil recruitment to the SI and found that germ-free (GF) and specific-pathogen-free (SPF) colonized BALB/c mice harbored similar eosinophil frequencies and numbers (Figure 1E), confirming that their initial seeding into the SI is independent of the microbiota (Mishra et al., 1999). Furthermore, eosinophil distribution within villi was not influenced by colonization status at steady state (Figure 1F). All together these data characterize eosinophil localization within the villous lamina propria and corroborate earlier findings of their microbiota-independent recruitment to the SI.

### Absence of resident SI eosinophils leads to villous abnormalities and alterations in the transcriptome

In order to further assess the potential contributions and importance of eosinophil residency in the SI, we analysed the impact of eosinophil-deficiency using the widely employed Δdbl.GATA1 mouse strain (Yu et al., 2002). As expected, eosinophils were absent in the SI of Δdbl.GATA1 mice (Figure 2A). Intriguingly, whole-mount imaging revealed that an absence of eosinophils impacted villous architecture characterized by a reduction in villous surface area, most pronounced in the proximal SI where eosinophils are most abdundant (Figure 2A). We also observed that NOD.SCID x γc^−/−^ mice, which have diminished SI eosinophil numbers (Figure S2A) (Carlens et al., 2009), exhibit a similar villous phenotype (Figure S2B). These villous defects were not simply due to reduced cellularity resulting from eosinophil loss as Rag2^−/−^ mice, which have drastically reduced immune cellularity due to loss of T and B cells, did not exhibit villous defects (Figure S2C). In order to confirm whether the villous alterations were eosinophil-specific and not due to an unappreciated impact of the Δdbl.GATA1 mutation in non-hematopoietic cells, we generated bone-marrow chimeras in which Δdbl.GATA1 bone marrow was transferred into irradiated BALB/c recipients. These data confirmed that Δdbl.GATA1 mutation within the hemopoetic compartment was responsible for the observed villous phenotype (Figure S2D). Despite alterations in villous architecture, Δdbl.GATA1 mice exhibited similar numbers of CD45^−^ stromal cell populations and normal proportions of the main stromal cell subsets including fibroblasts, lymphatic and blood endothelial cells (Figure S2E).

**Figure 2:**
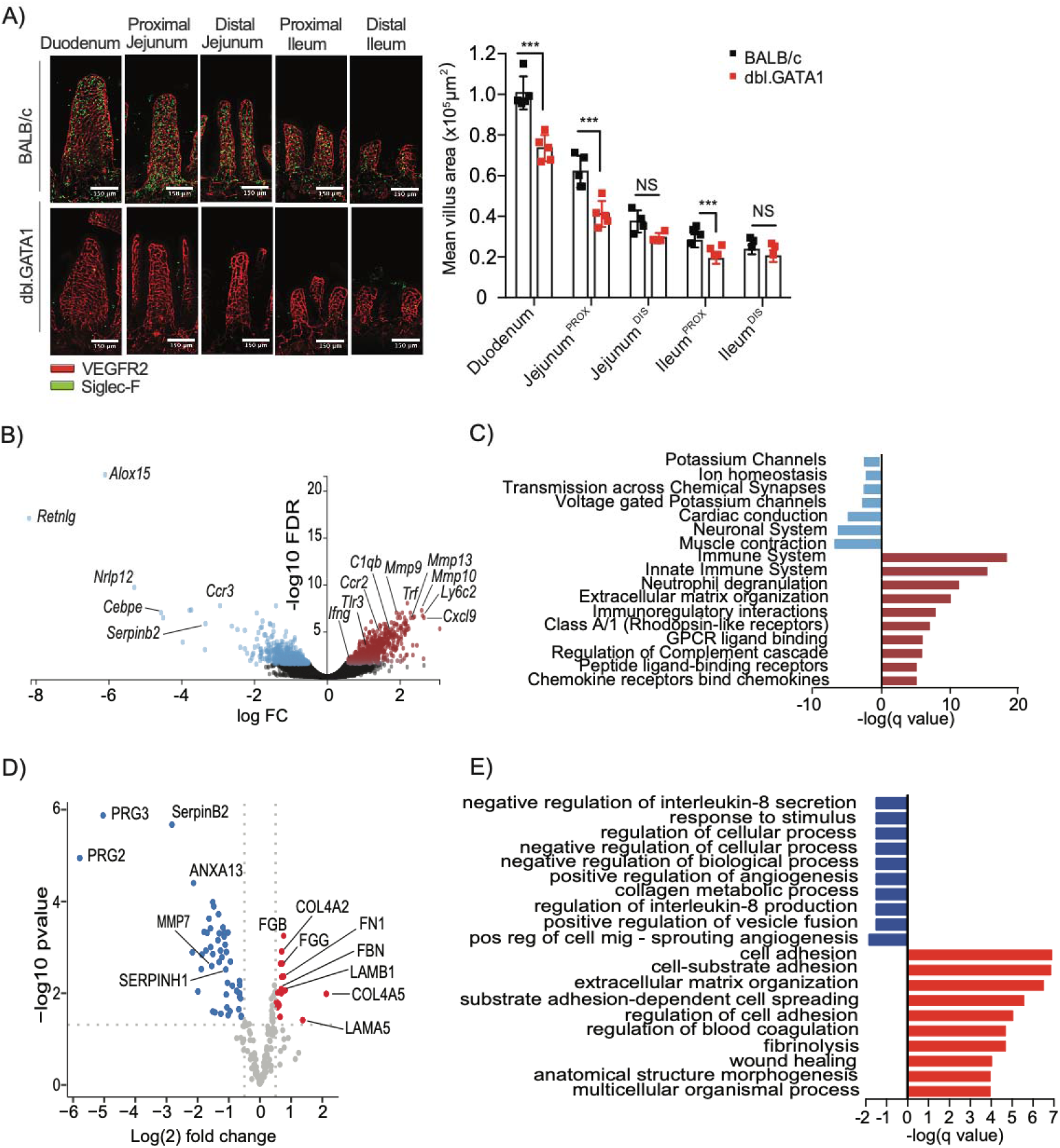
Eosinophil deficiency results in villous atrophy and is associated with transcriptome alterations within the small intestinal (SI) lamina propria. (A) Quantification of mean villous surface area from various SI regions calculated from the vascular cage area of whole-mount tissues from SPF BALB/c and Δdbl.GATA1 mice stained for eosinophils (Siglec-F, green) and blood endothelial cells (VEGFR2, red). Images show representative villi from a single mouse. The bar graph shows data from one experiment and are representative of at least 2 independent experiments performed using SPF mice from two different animal facilities. Statistical analysis was performed using a multiple t-test with statistical significance determined using the Holm-Sidak method; p<0.05*, p<0.01**, p<0.001***, N.S. non-significant, each bar graph represents data mean ± SD of n=5 mice per experiment with individual data points representing the mean surface area of at least >30 villi randomly sampled within the indicated area of tissue per mouse. (B-C) Bulk tissue RNA-seq of proximal jejunum, devoid of epithelial cells (non-IEC fraction), from BALB/c and Δdbl.GATA1 SPF mice: (B) Volcano plot showing all differentially expressed genes (DEGs) identified; (C) gene set enrichment analysis (GSEA) of gene pathways downregulated (blue) or upregulated (red) in ΔdblGATA1 mice (FDR <0.01, log2FC > 1). Only pathways with an interaction score of >0.7 are shown. (D-E) Tandem mass spectrometry (MS) on extracellular matrix (ECM) enriched proteins from the proximal jejunum of BALB/c and Δdbl.GATA1 SPF mice: (D) Volcano plot showing all proteins ECM and ECM-related proteins; (E) GSEA analyses of proteomic data showing pathways downregulated (blue) or upregulated (red) in Δdbl.GATA1 mice (FDR <0.05 and Log_2_FC>0.5) proteins in dbl.GATA1 mice. See also Figure S2.

To gain a broader perspective of the impact of eosinophil-deficiency within the SI, we analysed the transcriptome of intestinal epithelial cells (IEC+) and lamina propria (LP/IEC-) fractions by RNA sequencing (RNA-seq). Only one gene was significantly altered between the entire IEC transcriptome of BALB/c and Δdbl.GATA1 mice, suggesting that eosinophils do not influence epithelial stemness or differentiation (Figure S2F). In contrast, significant changes were observed in the transcriptome of the lamina propria, with 399 genes upregulated and 197 genes down regulated (FDR < 0.01 and Log2FC > 1) in the absence of eosinophils (Table S1). As expected, the most strongly downregulated transcripts were those known to be associated with eosinophils, including *Alox15* (Miyata et al., 2013; Uderhardt et al., 2017), *RetnIg* (Chumakov et al., 2004; Schinke et al., 2004), *Nlrp12* (Allen et al., 2012), *Cebpe* (Gombart et al., 2003), *Ccr3* (Pope et al., 2005) and *Serpinb2* (Fulkerson et al., 2006; Swartz et al., 2004) (Figure 2B). Gene Set Enrichment Analysis (GSEA) indicated that loss of eosinophils resulted in a positive enrichment of pathways related to innate immune/inflammatory responses, extracellular matrix (ECM) organization, G-protein-coupled receptors (GPCR) and GPCR ligand binding, and regulation of complement cascade (Figure 2C and Table S1). We also observed a negative enrichment for pathways related to muscle contraction and neuronal signalling (Figure 2C and Table S1).

The observed enrichment of transcripts from ECM-related pathways (Figure 2C) is in keeping with reported associations between eosinophils and tissue remodelling in the airways (Flood-Page et al., 2003; Humbles et al., 2004). This prompted us to investigate the impact of eosinophil deficiency on ECM composition in more depth by conducting quantitative mass spectrometry (LC-MS/MS) analysis of the ECM-enriched fraction of control BALB/c or Δdbl.GATA1 mice, using tissue obtained from the proximal jejunum where the villous phenotype is most pronounced. We observed significant changes in multiple ECM and ECM-related proteins, with 15 proteins increased and 42 proteins decreased (FDR <0.05 and Log_2_FC>0.5) in samples from Δdbl.GATA1 small intestine (Figure 2D and Table S2). The most downregulated proteins in Δdbl.GATA1mice included Serpin B2 and the Eosinophil Major Basic proteins (PRG2 and PRG3) (Figure 2D). Of note, eosinophil-deficiency decreased levels of MMP7, concomitant with increased amounts of Fibronectin and multiple subunits for the basement membrane proteins Collagen alpha-4(IV) and Laminin, which are all substrates for Matrix Metalloproteinease (MMP)-7 (Figure 2D) (McGuire et al., 2003). GSEA analyses of the proteomic data indicated pathways associated with cell and cell-substrate adhesion and the tissue damage response (coagulation, fibrinolysis) (Barker and Engler, 2017; Hubbard et al., 2014; Sottile and Hocking, 2002) were enhanced in dbl.GATA1 small intestine (Figure 2D and Table S2). Furthermore, eosinophil-deficiency slightly decreased the abundance of proteins involved multiple pathways, including those related to angiogenesis and IL-8 (Figure 2D and Table S2). Taken together, these data indicate that eosinophils impact villous gene expression and architecture, with an absence of these cells leading to increased villous atrophy associated with an increased transcriptomic signature of inflammation and proteomic evidence of tissue damage.

### Eosinophil deficient mice exhibit abnormalities in the intestinal barrier under homeostatic conditions

Given the alterations to tissue architecture, and the negative impact of eosinophil deficiency on angiogenic pathways, we assessed a possible role for eosinophils in regulating the villous vasculature by conducting a density and branching analysis of the blood vessels. However, no obvious defects with regards to vessel patterning were noted (Figure S3A). Many of the ECM proteins increased in the absence of eosinophils consituted specific components of the basement membrane which influences intestinal epithelial cell function (Groulx et al., 2011; Vllasaliu et al., 2014) and forms an anatomical barrier between the epithelium and underlying connective tissue. Thus we next investigated the impact of eosinophil deficiency on IEC homeostasis and barrier function. In agreement with our finding that IECs from WT and Δdbl.GATA1 mice did not exhibit significant alterations in gene transcription, the differentiation of IEC into Goblet cells, Tuft cells or Paneth cells was not altered between BALB/c and Δdbl.GATA1 mice (Figure S3B-D). IEC proliferation was also not affected by the absence of eosinophils (Figure 3A, Figure S3E). However, we did observe a moderate, but consistent, delay in IEC migration in Δdbl.GATA1 mice, as determined by tracking the distance travelled by EdU^+^ IECs from villus-crypt axis at 24h post-EdU labeling (Figure 3B). Delayed IEC migration was confirmed in bone marrow transfer experiments where transfer of Δdbl.GATA1 bone marrow to irradiated BALB/c recipients was sufficient to induce the same delay in IEC migration (Figure 3C). Intraepithelial lymphocytes (IELs) are interspersed between IECs above the basement membrane and reinforce barrier function by scanning for infected or stressed IEC (Cheroutre et al., 2011; Roberts et al., 1996). In line with alterations in IEC homeostasis, we also observed that Δdbl.GATA1 mice harbored decreased frequencies and numbers of CD8αα^+^ and CD8αβ^+^ TCRαβ^+^ IELs (Figure 3D).

**Figure 3.**
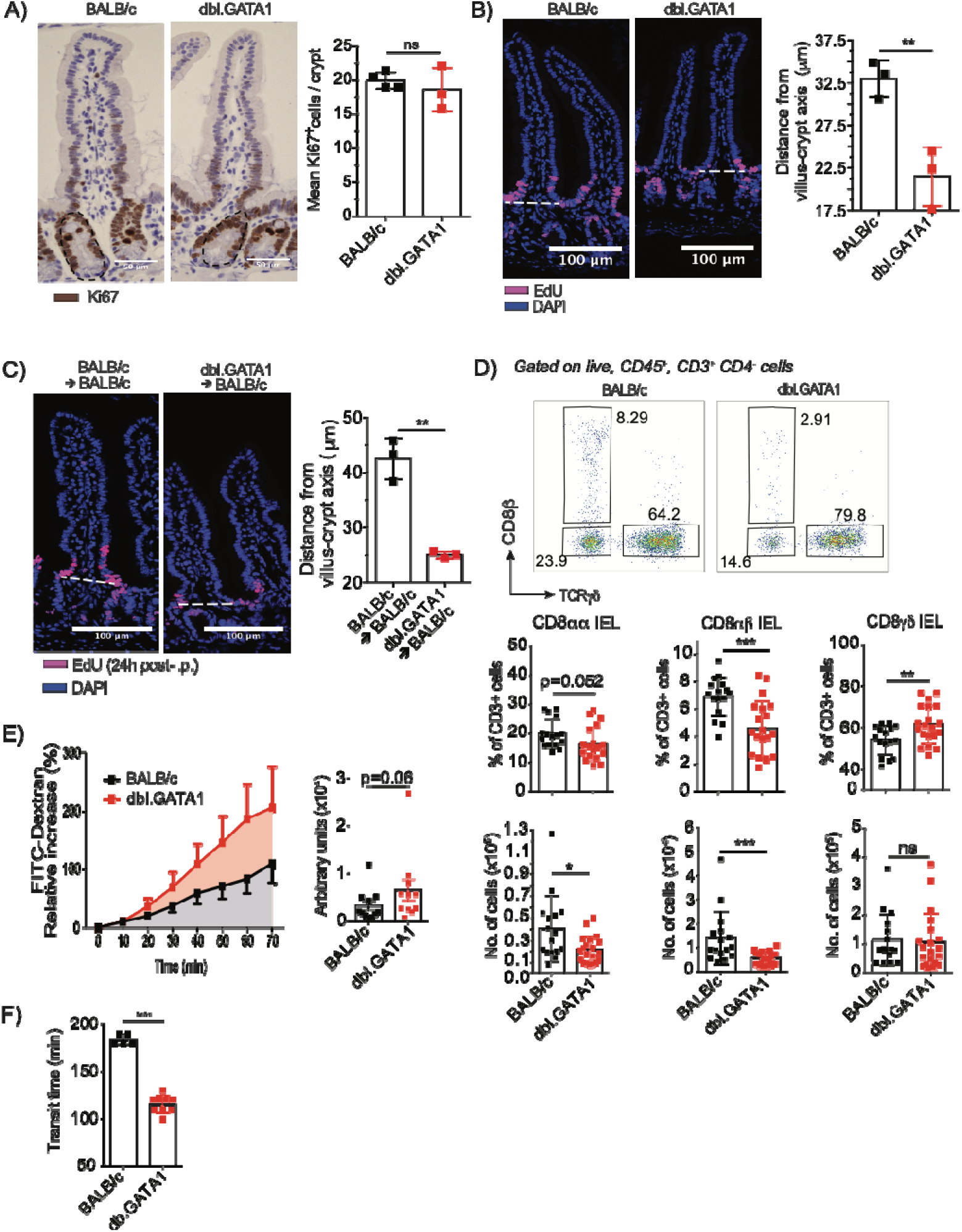
Eosinophil-deficient mice exhibit significant alterations to IEL populations, barrier integrity and gastro-intestinal motility. (A) Tissue sections from paraffin embedded jejunal tissues from BALB/c and Δdbl.GATA1 SPF mice stained using antibodies directed against Ki67, then counterstained with hematoxylin. Epithelial turnover was assessed by quantifying the average number of Ki67^+^ epithelial cells per crypt (dotted lines within representative image demarcating crypts) with each data point indicated as the average per mouse of at least >20 randomly sampled crypts. (B-C) Jejunal tissues were collected 24 hrs after intraperitoneal injection of EdU (200μg/mouse) injection. (B) Epithelial cell migration was assessed by measuring the distance of EdU^+^ cells migrated above the villus-crypt axis (indicated by dotted-line; EdU^+^ cells (purple), nuclei stained with DAPI (blue)). Panels show tissues from (B) BALB/c and Δdbl.GATA1 SPF mice, and (C) irradiated BALB/c recipients 8 weeks after bone marrow transfer of either BALB/c or Δdbl.GATA1. For panels, A-C each bar represents data mean ± SD of n=3 mice with individual data points the mean surface area of at least >30 villi randomly sampled within the jejunum tissue per mouse (D) Representative FACS plots showing gating strategy for CD8^+^ intraepithelial lymphocytes (IELs) subpopulations from BALB/c and Δdbl.GATA1 SPF mice. Bar graphs showing frequency of CD8^+^ IELs within live CD45^+^ CD3^+^ CD4^−^ cells and absolute cell numbers. Each bar represents data mean ± SD of n=15-20 mice with individual data points representing a single mouse. (E) Barrier integrity was analysed using proximal jejunum fractions (2 cm) from BALB/c and Δdbl.GATA1 SPF mice, with paracellular permeability to FITC-Dextran (4 kDa) evaluated every 10 minutes for 70 minutes using a Ussing Chamber. Bar graph shows area under the curve. (E) Total intestinal transit time in BALB/c and Δdbl.GATA1 SPF mice was assessed by oral gavage of Carmine Red (180 mg/mL) followed by faecal pellet harvesting. Barrier integrity and transit time plots are pooled from 2-3 independent experiments (n= 5-11) with individual data points each representing a single mouse. Error bars indicate SD. For all panels statistical analysis was performed using a student t-test p<0.05*, p<0.01**, p<0.001***, N.S. non-significant. See also Figure S3.

We next investigated whether the observed changes in basement membrane proteins, defects in epithelial cell migration and altered IEL frequencies affected barrier function in Δdbl.GATA1 mice. We observed that *ex vivo* explants of the small intestine of Δdbl.GATA1 mice exhibited a slight increase in barrier leakage (Figure 3E). This increase in barrier leakage may explain the increased inflammatory signature (Figure 2C and Table S1), that was associated with a predicted activation of regulators associated with bacterial (LPS, PolyI:C) or inflammatory pathways (IFNγ, TNF, IFNα, IRF3, MyD88, STAT1) (Table S3). These findings suggest that eosinophils may reinforce villous integrity by regulating ECM turnover and preventing activation of inflammatory responses by intestinal microbes.

In contrast to the positively enriched ECM signature in the lamina propria RNA-seq data, negative enrichment for neuronal and muscle contraction pathways was observed in Δdbl.GATA1 mice (Figure 2C). Given the proximity of eosinophils to neuronal/glial cells of the enteric nervous system (ENS) (Figure 1C, Figure S1B), and the importance of the ENS in regulating gastrointestinal motility, we evaluated whether eosinophil-deficiency impacted on this critical physiological function. Based on the migration of carmine red dye (Dey et al., 2015), Δdbl.GATA1 mice had faster total intestinal transit time (Figure 3F). Altered transit correlated with changes in muscle contractility with Δdbl.GATA1 mice exhibiting a significant increase in the average cyclic height of contractions of SI sections taken from the upper SI and placed in a longitudinal orientation under 1 g tension in an organ bath (Figure S3F). A trend towards decreased contraction frequency was also observed in the uppermost part of the SI from Δdbl.GATA1 mice (p<0.0566), whilst the tension remained similar over time between both Δdbl.GATA1 and BALB/c mice (Figure S3F). Collectively, these results indicate that tissue-resident eosinophils influence various aspects of SI physiology during homeostasis, including villous architecture, barrier maintenance and intestinal motiility. Such observations provide insight into the functional relevance for the residence of eosinophils within the small intestine.

### Microbial colonization drives homeostatic SI eosinophil function

Given the recruitment and maintenance of eosinophils to the SI prior to microbiota colonization, and the barrier and physiological defects that result from eosinophil loss in SPF mice, we hypothesized that eosinophils may be important for mitigating microbiota-induced inflammation and alterations to intestinal function. In support of this, we observed that MyD88^−/−^ mice, which lack microbial sensing (Kawai et al., 1999), exhibited a similar villous defect to Δdbl.GATA1 mice (Figure S4A). To address whether the intestinal microbiota influences eosinophil-mediated maintenance of the intestinal barrier, we re-derived Δdbl.GATA1 mice to germ-free (GF) status. Strikingly, Δdbl.GATA1 and control BALB/c mice exibited similar villous surface areas when reared under GF conditions (Figure 4A). In contrast, microbial colonisation resulted in marked villous atrophy in both Δdbl.GATA1 and BALB/c mice, but the presence of eosinophils in BALB/c mice mitigated the extent of the induced atrophy (Figure 4A). This observation was not due to alterations in cecal microbial community composition as no changes in bacterial family members were observed (Figure 4B). As the CD8αβ^+^TCRαβ^+^ IEL populations that were decreased in Δdbl.GATA1 SPF mice are known to expand in a microbiota-dependent manner (Chen et al., 2018; Williams et al., 2004), we analyzed CD8αβ^+^TCRαβ^+^ IEL in GF and re-colonized Δdbl.GATA1 mice. As expected, re-colonization of control BALB/c GF mice resulted in specific expansion of this IEL population (Figure 4C). In contrast, the frequency of CD8αβ^+^TCRαβ^+^ IEL in the SI of GF Δdbl.GATA1 mice did not increase following microbial colonization, indicating that eosinophils are required for this process (Figure 4C). In line with these findings, we also observed that GF Δdbl.GATA1 and BALB/c mice did not exhibit differences in permeability or intestinal transit time (Figure 4D and 4E). Microbial colonization resulted in increased permeability and decreased transit time, and in both cases these changes were amplified in the absence of eosinophils (Figure 4D and 4E).

**Figure 4.**
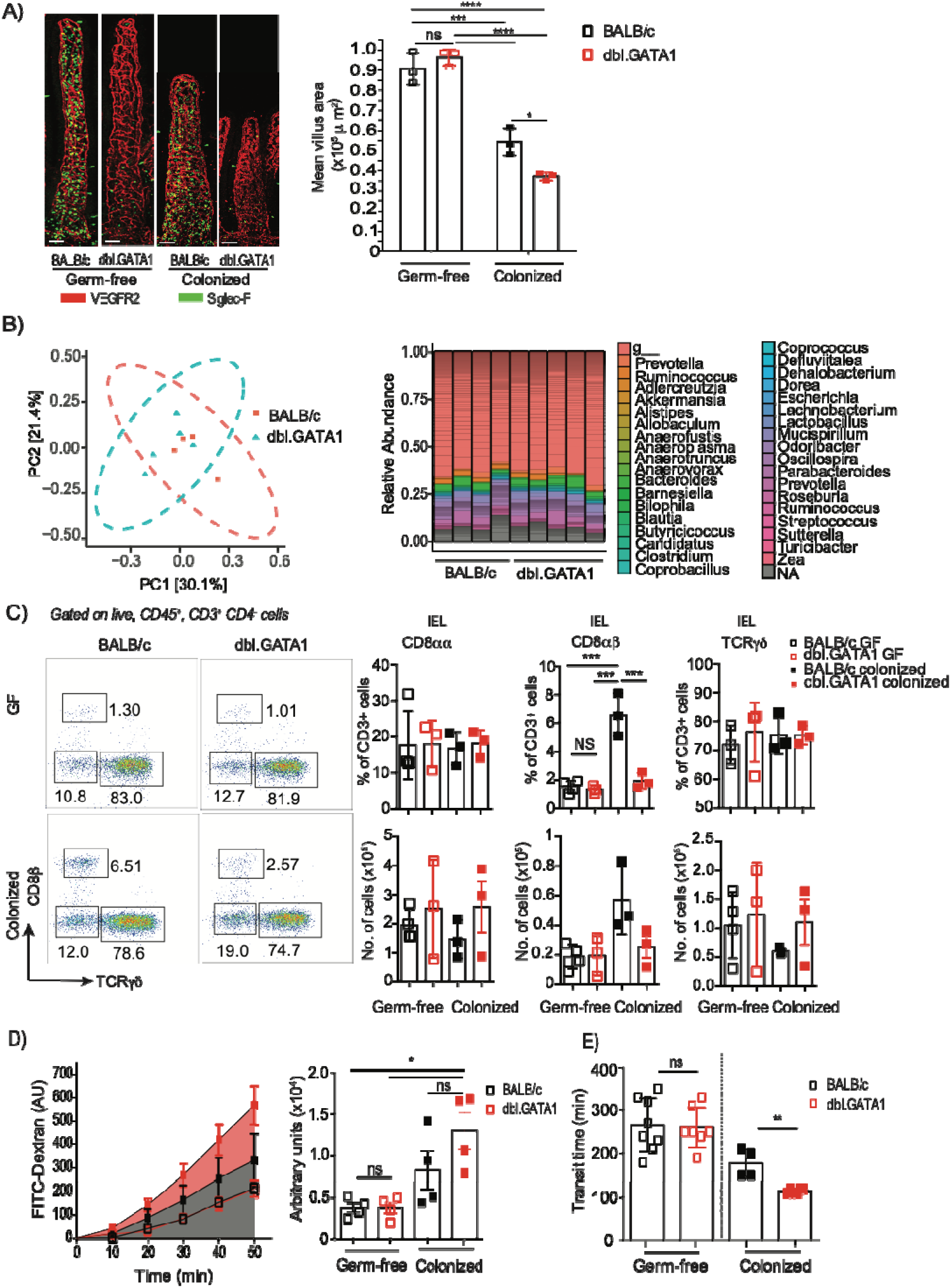
Eosinophils regulate microbial-induced villous size alterations, epithelial barrier integrity and intestinal transit. Germ-free (GF) BALB/c and GF eosinophil deficient (Δdbl.GATA1) mice were compared to previously germ-free (ex-GF) BALB/c and GF Δdbl.GATA1 mice recolonized for 4 weeks with microbiota from specific-pathogen free (SPF) BALB/c donors. (A) Whole-mount tissues from the proximal jejunum were stained for eosinophils (Siglec-F, green) and blood endothelial cells (VEGFR2, red). Images show representative villi from a single mouse. Bar graphs show quantification of mean villous surface area, calculated using the vascular cage. Data from one experiment are shown and indicate mean ± SD of n=3 mice with individual data points representing the mean surface area of at least >30 villi randomly sampled within the jejunum tissue per mouse. Data are representative of 2 independent experiments. Statistical analysis was performed using two-way ANOVA, Tukey’s multiple comparisons post-test; p<0.05*, p<0.01***, p<0.0001****, n.s. non-significant. (B) 16S sequencing analysis of cecal contents harvested from recolonized BALB/c and GF Δdbl.GATA1 mice. (C) Representative FACS plots showing gating strategy for CD8^+^ intraepithelial lymphocytes (IELs) subpopulations from BALB/c and Δdbl.GATA1 mice. Bar graphs showing frequency of CD8^+^ IELs within live CD45^+^ CD3^+^ CD4^−^ cells and absolute cell numbers. Statistical analysis was performed using two-way ANOVA, Tukey’s multiple comparisons post-test; p<0.05*, p<0.01**, p<0.001***, N.S. non-significant, each bar graph represents data mean ± SD of n=3-4 mice with individual data points representing a single mouse. Data are representative of 5 independent experiments, where 3 out 5 experiments showed significant differences between BALB/c and dbl.GATA1 recolonized mice. (D) Barrier integrity was analysed using proximal jejunum fractions (2 cm). Paracellular permeability to FITC-Dextran (4 kDa) was evaluated over a total period of 70 minutes using an Ussing Chamber. Bar graph shows area under the curve. (E) Total transit time was assessed by oral gavage of Carmine Red followed by faecal pellet harvesting. Barrier integrity and transit time plots are representative of 2-3 independent experiments (n=4-8 mice) with individual data points representing a single mouse. (C-D) Statistical analysis were performed using a two-way ANOVA, Tukey’s multiple comparisons post-test or (E) student t-test; p<0.05*, p<0.01**, p<0.001***, N.S. non-significant. See also Figure S4.

Bacterial translocation to extra-intestinal tissues can occur during homeostasis and lead to the induction of systemic anti-commensal immunoglobulin (Ig)-G (Zeng et al., 2016). To validate a role for eosinophils in regulating barrier integrity *in vivo*, we monocolonized GF Δdbl.GATA1 and BALB/c mice with *Akkermansia muciniphila*, a known inducer of IgG-class switching (Ansaldo et al., 2019), and determined *A. muciniphila* specific IgG levels in the serum using bacterial flow cytometry. In support of the observed increase in intestinal permeability observed in following complex microbial colonization (Figure 4D), monocolonized Δdbl.GATA1 mice displayed signficantly increased serum levels of bacterial-specific IgG2a, IgG2b and IgG3 compared to monocolonized BALB/c mice (Figure S4B). We then assessed whether the ECM-alterations observed in SPF Δdbl.GATA1 mice (Figure 2E) were also driven in response to the microbiota. Proteome analysis of whole jejunum tissue segments from GF and recolonized Δdbl.GATA1 and BALB/c mice revealed no significant changes in ECM or ECM affiliated proteins between GF Δdbl.GATA1 and BALB/c mice (Table S4). In contrast, and in line with our findings in SPF mice, recolonization of GF Δdbl.GATA1 and BALB/c mice led to increases in a number of ECM proteins, including laminins and tight /adherens junction-related proteins, which was exacerbated in the absence of eosinophils (Figure S4C and Table S5). These data suggest that alterations in the ECM in the absence of eosinophils are driven in response to the microbiota. Collectively, these observations indicate that eosinophils respond to microbial colonization and this response is required to promote SI tissue homeostasis.

### The microbiota is a key regulator of eosinophil activation and heterogeneity in the SI

To determine whether eosinophils respond directly to microbial colonization, we investigated the turnover of SI resident eosinophils in GF versus recolonized GF Δdbl.GATA1 and BALB/c mice in a BrdU pulse-chase experiment. Microbial colonization resulted in significantly shorter half-life of intestinal eosinophils, consistant with increased cellular activity and reduced survival (Figure 5A). Microbial colonization likely led to increased eosinophil degranulation, as suggested by the presence of sombrero vesicles and empty vesicles in SPF eosinophils as determined by transmission electron microscopy (TEM; Figure 5B), and lower levels of intracellular eosinophil peroxidase (EPO; Figure 5C). These data suggest that homeostatic degranulation of SI eosinophils is driven by the intestinal microbiota. To further investigate microbial-induced eosinophil activation, we conducted mass cytometry (CyTOF) analysis of SI eosinophils isolated from GF and recolonized BALB/c mice. Unsupervised analysis of the surface expression of CD11c, Siglec F, CD11b and F4/80 on eosinophils by CyTOF identified four clusters of eosinophils in the SI of GF and re-colonized mice (Figure 5D). Microbial colonization resulted in a reduced frequency of clusters characterized by lower (#1) to intermediate (#2 and #3) expression levels of CD11b, Siglec-F and F4/80, whilst cluster #4, distinguished by higher expression levels of all four markers, increased (Figure 5D). Microbial colonization also led to an increase in the expression of Gr1 on clusters 1, 2 and 3, an increase in CCR3 expression in cluster 3, and an increase in ST2 expression in cluster 2 (Figure 5E). Given the previous association of these makers with eosinophil activation in response to intestinal inflammation (Arnold et al., 2018; Griseri et al., 2015), these data indicate that microbial colonization can regulate the activation and/or maturation of intestinal eosinophils. Taken together, our results show that eosinophils respond to microbial colonization and play a critical role in facilitating homeostatic host-microbiota crosstalk by maintaining barrier integrity and regulating microbiota-dependent changes in intestinal physiology.

**Figure 5.**
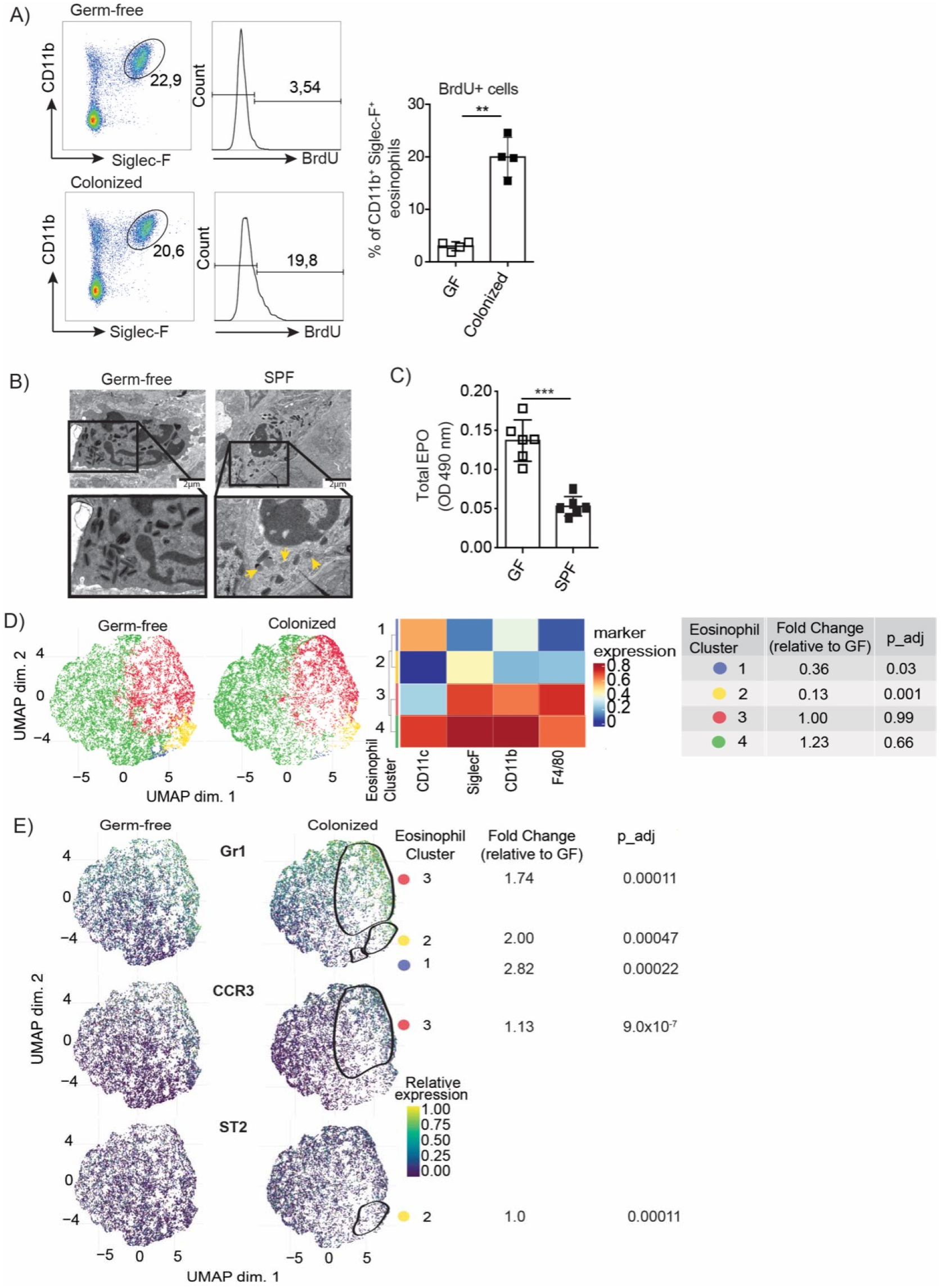
The intestinal microbiota modulates the activation of small intestine (SI) eosinophils. (A) SI eosinophils were analysed in GF and SPF BALB/c mice after all mice received BrdU in the drinking water for 8 days. Plots are representative of 2 independent experiments (n=2-4 mice) (B) Transmission electron microscopy (TEM) SI eosinophils from GF and SPF BALB/c mice, the yellow arrows in the SPF eosinophil indicating empty vesicles. (C) Total eosinophil peroxidase (EPO) activity were analysed in FACS-sorted eosinophils from the SI of GF and SPF BALB/c mice. Plots are pooled from 3 independent experiments (n=2). Individual data points represent a pooled of 8-10 mice for the acquisition of 1×10^5^ sorted eosinophils/well. (D, E) CyTOF analysis of proximal jejunum eosinophils from GF and SPF BALB/c mice (n=3-4). Cells were manually pre-gated on singlets (191IrDNA-event length), live (195Pt cisplatin-), 89Y CD45+ cells on FlowJo and individual FSC files were concatenated for each group and exported for analysis by unsupervised method (D) Proximal jejunum eosinophils were identified and clustered using uniform manifold approximation and projection (UMAP) algorithm. Heat map of surface marker expression considered for eosinophil identification and cluster analyses showed in the UMAP representation. Eosinophils numbers on each cluster were determined and significant differences within clusters are represented as fold change relative to GF (E) Differential expression of Gr1, CCR3 and ST2 on within eosinophil cluster represented as fold change relative to GF in the UMAP representation.

## DISCUSSION

In addition to the role of the microbiota in shaping immune cell repertoire, the intestine undergoes significant morphological and physiological alterations, with GF mice exhibiting thinner and longer villi (Smith et al., 2007), decreased permeability (Hayes et al., 2018), and reduced intestinal motility (Muller et al., 2014), when compared to mice colonized by a complex microbiota. The finding that eosinophil-deficient mice exhibit exaggerated microbial-induced villous atrophy and increased intestinal permeability following microbial colonization reveals an unexpected function for these cells as crucial regulators of the host adaptation to the presence of intestinal bacteria. Luminal bacterial products are known to induce inflammation and an associated tissue remodelling response (van Tol et al., 1999), and our RNAseq data revealed that eosinophil-deficiency resulted in the upregulation of inflammatory pathways including enrichment of neutrophil degranulation and complement cascade genes. Thus our observations that eosinophils modulate microbial-induced alterations to intestinal morphology and permeability, alongside the presence of a ECM signature indicative of persistent tissue remodelling in the absence of eosinophils, may well result from a role for these cells in dampening microbial-induced inflammation. Mechanistically, eosinophils may regulate intestinal integrity by directly contributing to the tissue remodelling processes, either via alterations to fibroblast function or direct interactions with the ECM. The latter hypothesis is supported by the abiltiy of eosinophils to secrete an array of products with the potential to modulate ECM deposition and degradation (Shah et al., 2020), and with reports linking eosinophil function to tissue remodelling in the breast (Gouon-Evans et al., 2002; Gouon-Evans et al., 2000) and uterus (Vicetti Miguel et al., 2017). The increased intestinal permeability apparent in eosinophil-deficient Δdbl.GATA1 mice may also be linked to the delayed epithelial cell migration and associated alterations in basement membrane proteins noted. The basement membrane plays a crucial role in guiding the migration of newly differentiated enterocytes from the crypt to villous tip (Heath, 1996). Of note, we observed increases in ECM components that drive epithelial migration and re-epithelialization/wound closure (e.g. Collagen VI, Laminins) and a decrease in proteases known to regulate the turnover and/or structure/bundling of these components, including MMP7, Cathepsin B and Serpin H1 (Li et al., 2016). These findings suggest that SI eosinophils may function as both positive and negative regulators of ECM deposition organization and composition under homeostatic conditions. Alterations in the ECM, particularly enzymes differentially expressed either at a transcriptomic and/or proteomic level, are also noted to influence the bioavailability of growth and migratory factors, which could have far reaching effects on immunity and regenerative responses (Chester and Brown, 2017). Thus, altered ECM composition may also contribute to the observed requirement for eosinophils in maintaining microbial-induced SI IEL populations. Reduced IEL numbers in Δdbl.GATA1 mice could in turn contribute to increased intestinal permeability. Our findings that eosinophils promote barrier integrity following microbial colonisation are in keeping with previous studies reporting a requirement for these cells in barrier integrity and pathogen control following infection with *Clostridium difficile* (Buonomo et al., 2016).

In addition to barrier and villous defects, we also provide evidence for SI eosinophils in influencing the ENS. Eosinophils were found in close contact with enteric glia and neural processes which extend through the villous, and our transcriptomic analysis of Δdbl.GATA1 mice revealed significant alterations in pathways related to neuronal and muscular signaling. These findings were associated with alterations in the patterning of cyclic muscle contractions and increased intestinal motility under steady-state conditions. Of note, various receptors (e.g. *Adrb3*, *Adcyap1r1*, *Gfra1*), adhesion molecules (e.g. *Nrxn2/3*) and potassium/calcium channels (e.g. *Kcnj12/2/3*, *Cacna1b/c*) involved in the regulation of neuronal action-potential/excitability were differentially expressed in Δdbl.GATA1 mice. As our findings are based on bulk-RNAseq, it is difficult to gage the precise influence of eosinophils in the regulation of the ENS, and it is also possible these alterations related directly to impacts on muscle cells. However, given the close proximity of eosinophils to neuronal axons, and past reports linking eosinophils to altered neuronal growth function (Drake et al., 2018), it is likely that eosinophils can regulate aspects of ENS function and the data presented here warrants further, more focused, investigations of eosinophil-ENS interactions.

Microenvironmental cues specific to the intestine have been proposed to regulate eosinophil survival (Carlens et al., 2009; Mishra et al., 1999). To date, specific factors shown to be important include eotaxin (Mishra et al., 1999) and γc-dependent signals (which include IL-4, IL-7, IL-15 and IL-21) (Carlens et al., 2009). Our results reveal the microbiota as an additional regulator of eosinophil turnover within the small intestine. Although the recruitment of eosinophils does not require the microbiota, microbial colonization clearly impacts these cells, resulting in increased turnover and an altered activation state. Our data indicates that the intestine contains at least 4 different clusters, or subpopulations, of eosinophils as defined by their surface phenotype. This is in keeping with a number of recent studies suggesting that eosinophils can exist as a heterogeneous group of cells, whose biology and phenotype is modulated by environment cues (Abdala-Valencia et al., 2018; Shah et al., 2020). Xenakis *et al* identified an intestinal CD11c^high^ eosinophil subpopulation that was closely associated with the IEL fraction and distinct from eosinophils found in the intestinal lamina propria and blood (Xenakis et al., 2018). In models of allergic airway inflammation eosinophils upregulate levels of Siglec⌞F and CD11c and exhibit altered morphology, including increased vacuolarization and segmentation of the nucleus, following their migration from the lung interstitium into the airways (Abdala Valencia et al., 2016). In a second study, lung resident eosinophils exhibiting intermediate levels of Siglec-F were described to exert regulatory functions, whilst inflammatory eosinophils recruited in response to allergen challenge were defined by increased Siglec-F expression and a more segmented nucleus. These cells were also able to migrate into the airways and contributed to inflammation (Mesnil et al., 2016). The intestine is unique in that it is constantly subject to a high level of remodelling and epithelial cell turnover, which is driven by the presence of a complex microbiota (Agace and McCoy, 2017). Thus, it is perhaps not surprising that microbial colonization leads to a specific increase the eosinophil cluster defined by the highest expression levels of Siglec⌞F and CD11c, akin to that observed in response to inflammatory subsets within the lung. Further studies employing refined techniques (i.e. scRNA-seq) will be necessary to understand whether phenotypic changes observed in intestinal eosinophils in response to microbial colonization correlate with specific cell functionality/behaviour, and how each cluster is affected during infection or other inflammatory insults. Moreover, it remains unclear whether the microbiota may be regulating eosinophil heterogeneity and activation through direct or indirect mechanisms. It is possible that the physiological changes induced by microbial colonization induce eosinophil activation through tissue stress and damage signals (Smith et al., 2007). Alternatively, eosinophils are noted to express pattern recognition receptors (PRR) (Nagase et al., 2003) potentially allowing direct bacterial recognition and consequent eosinophil activation.

In summary, our findings demonstrate a key role for eosinophils in facilitating host adaptation to microbial colonisation by regulating microbial-induced alterations to intestinal morphology and physiology. This likely involves concerted interactions with the ECM and with other key cellular nodes within the villous lamina propria. Furthermore, we reveal intestinal eosinophils to be a heterogeneous population whose phenotype is modulated by microbiota-driven signals. These signals, either directly or indirectly, act as a key regulator of eosinophil activation and maturation within the intestine, thus arming these cells with the ability to modulate host tissue function. Our findings that eosinophils impact on key structual compentents of the intestine, including barrier integrity, ECM organization and ENS function, widen our appreciation of how these cells influence the function of healthy tissues. Our work indicates that a more holistic evaulation of tissue function is needed when assessing the mechanisms underlying eosinophil function in both health and disease. Furthermore, we hope our work will warrant further investigation into the mechanisms underlying microbial-, or inflammation-, driven activation of distinct eosinophil subsets. Understanding these mechanisms could potentially provide insight into the factors that underlie the nature of intestinal homeostasis versus disease and may provide novel therapeutic targets for the intervention of eosinophil-associated disorders.

## Author Contributions

KS, AI, JB-L, YK, GC, MM, RH, NCW, CAT, RB, and AD performed experiments and analyzed data. FA and HR performed bioinformatics. KS, AI, NLH and KDM wrote the manuscript, and all authors revised the manuscript and approved its final version. NLH and KDM conceived the project. TVP, NLH and KDM supervised the project.

## Acknowledgments

We thank M. Dicay for performing the Ussing chamber experiments. AI was supported by a Fellowship from São Paulo Research Foundation (2016/15882-7) and an Eyes High Postdoctoral Fellowship from the University of Calgary. This work was supported by grants from the Swiss National Science Foundation (SNSF310030_134902 to KDM, SNSF310030_156517 to NLH, and 31003A-156266 to TVP) and the European Research Council (ERC, FP/2007-2013) Agreement no. 281785 to KDM. KDM is also supported by a Canadian Institutes of Health Research (CIHR) grant (PJT-165930), a Canadian Foundation for Innovation (CFI) John R. Evans Leaders Fund (JELF) grant and the Cumming Medical Research Fund (CMRF). NLH is supported by a National Health and Medical Research Council (NHMRC) of Australia SRF-B fellowship. AD was supported by an NSERC Discovery Grant (DGECR-2019-00112). The IMC is supported by the Cumming School of Medicine, University of Calgary, Western Economic Diversification (WED) and Alberta Economic Development and Trade (AEDT), Canada.

## Supplementary Information

**Supplementary Figure 1.**
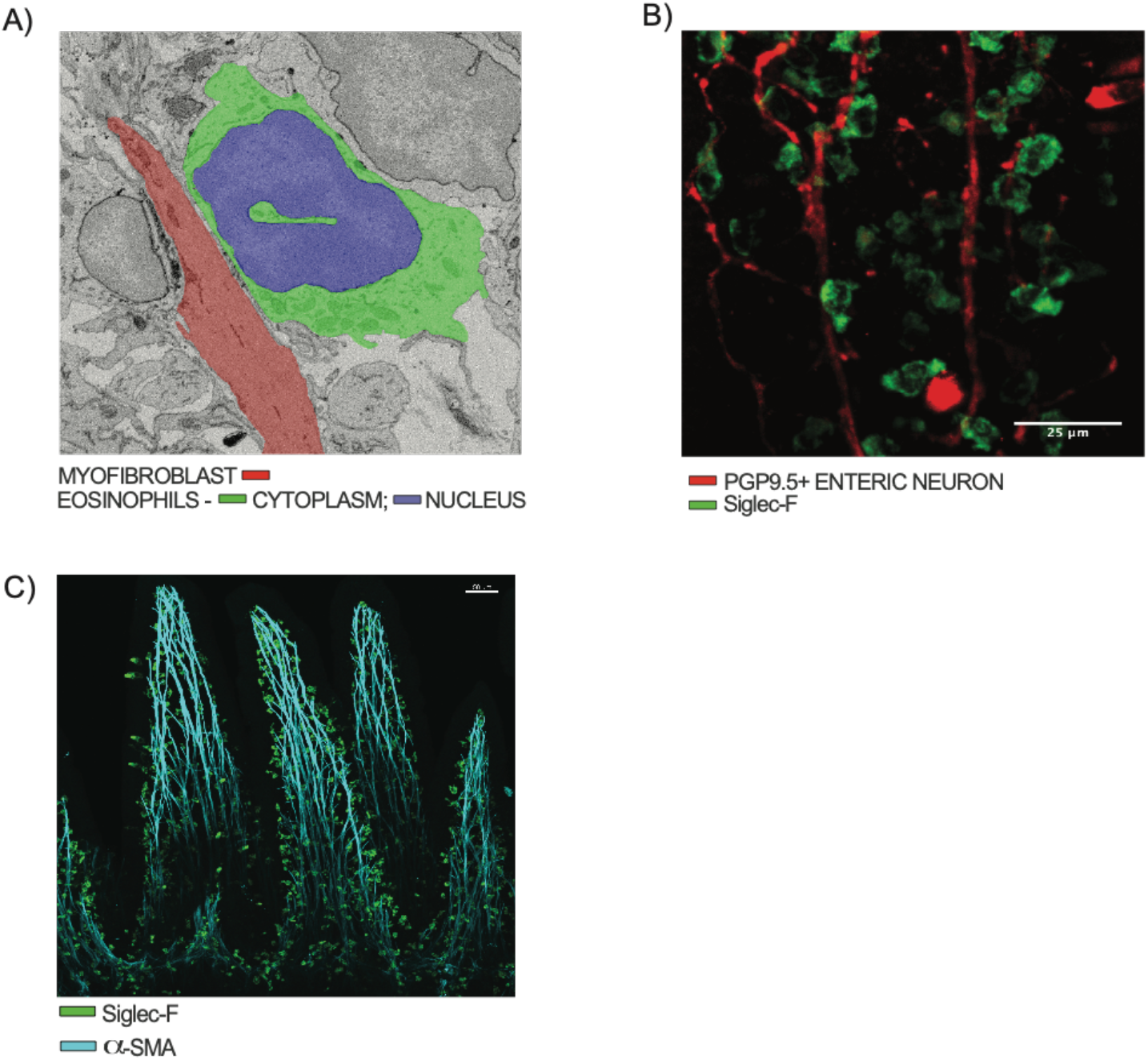
SI eosinophils are in close contact with structural cells within the lamina propria. Transmission Electron microscopy image of jejunum from specific-pathogen-free (SPF) BALB/c mice showing an eosinophil (green) in direct contact with a myofibroblast (red); digital colouring was used to highlight cells. (B) Whole mount image of jejunum from SPF BALB/c mice showing Siglec-F+ eosinophils (green) interacting with PGP9.5+ enteric neuronal axons (red) within the villous lamina propria. (C) Whole mount image of jejunum from SPF BALB/c mice showing α-SMA+ myofibroblasts (blue) and Siglec-F+ eosinophils (green) throughout the villi and crypt regions. Related to Figure 1.

**Supplementary Figure 2.**
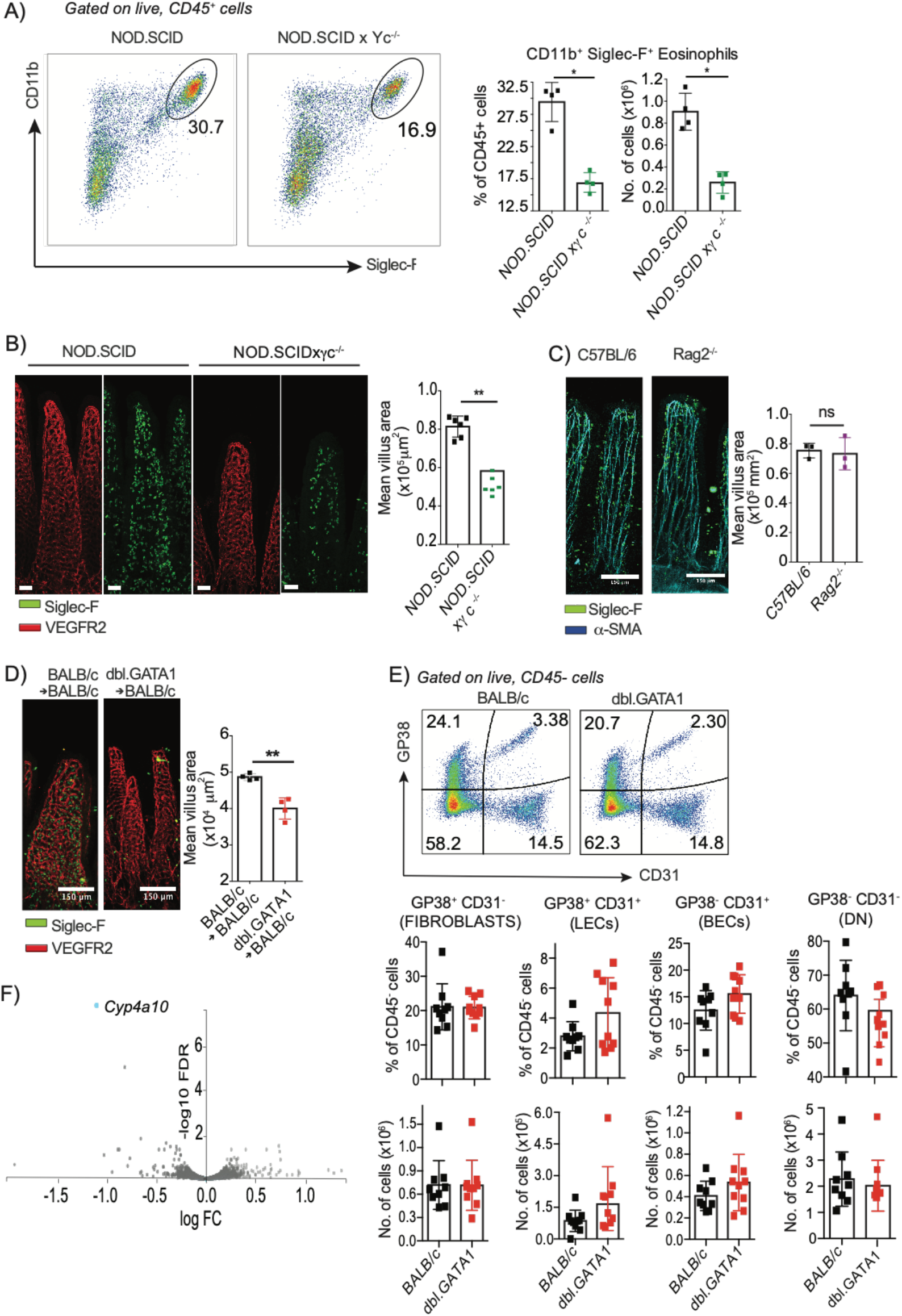
Villous atrophy in eosinophil-deficient mice is not due to reduced cellularity. (A) Flow cytometry analysis of eosinophils isolated from the SI lamina propria showing gating strategy, frequency of eosinophils within total CD45+ cells and absolute eosinophil numbers in the small intestinal lamina propria of NOD.SCID vs. NOD.SCIDγc^−/−^ SPF mice. (B-C) Quantification of mean jejunal surface area calculated using whole-mount tissues from (B) NOD.SCID vs. NOD.SCIDγc^−/−^ SPF mice (C) C57BL/6 vs. Rag2^−/−^ SPF and (D) irradiated BALB/c recipients 8 weeks after bone marrow (BM) transfer of BALB/c or Δdbl.GATA1 mice. Villous surface area (B-D) was calculated from the vascular cage following staining for eosinophils (Siglec-F, green) and blood endothelial cells (VEGFR2, red) or (C) eosinophils (Siglec-F, green) and smooth muscle fibers (α-SMA, cyan) (E) Flow cytometry analysis of gp38 and CD31 expression by stromal cells isolated from jejunum of BALB/c and μdbl.GATA1 mice indicating the frequencies and numbers of fibroblasts, lymphatic endothelial cells (LECs), blood endothelial cells (BECs), and double negative (DN) cells. Statistical analysis was performed using a student t-test; p<0.05*, p<0.01**, p<0.001***, N.S. non-significant, each bar graph represents data mean ± SD of n=3-10 mice with individual data points representing a single mouse (A&E) or the mean of 100 villi counted within the indicated area of tissue from a single mouse (B-D). (F) Volcano plot showing all genes identified from RNA-seq of epithelial cells enriched from the proximal jejunum (IEC fraction) of BALB/c and Δdbl.GATA1 SPF mice. Related to Figure 2, Table S1, Table S2.

**Supplementary Figure 3.**
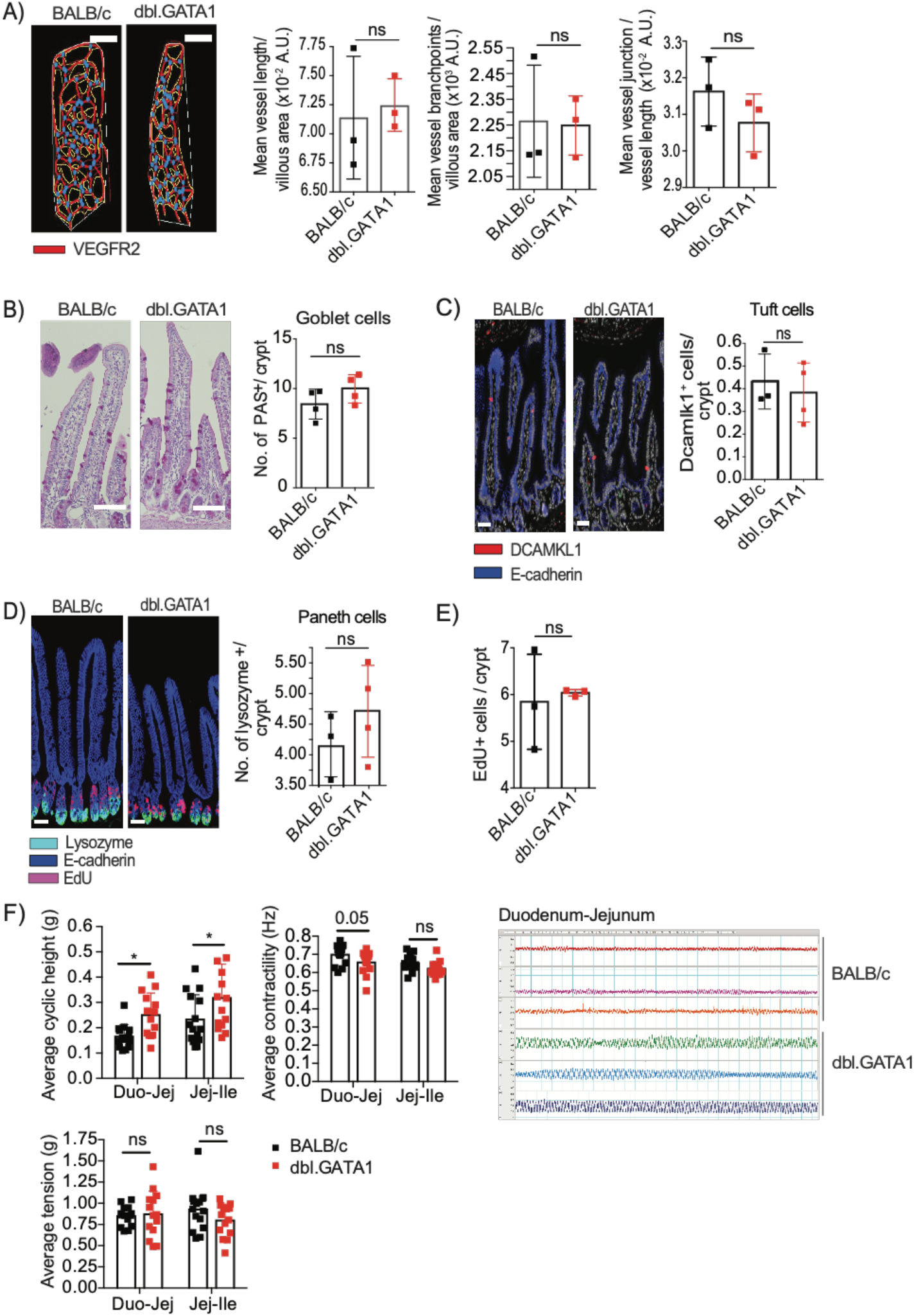
Eosinophil-deficient (Δdbl.GATA1) mice exhibit normal IEC differentiation but altered IEC migration and muscular contractility. (A) Whole-mount staining of jejunum from BALB/c and μ dbl.GATA1 SPF mice were stained for blood endothelial cells (VEGFR2, red). Blood vessels (VEGFR2+) were traced from villi images and ran on angiotool software – readouts were average number of junctions and branch points in the vessel network, in addition to vessel length. Images show representative images of traced vessel networks (white line), transposed onto the original image. Bar graphs show data normalized to villous surface area. Each data point represents a mean of >15 villi per mouse (n=3). (B-E) Tissue sections from paraffin embedded jejunal tissues from BALB/c and Δdbl.GATA1 SPF mice were stained with the indicated markers. Tissues were stained with antibodies against (B) goblet cells (periodic-acid Schiff; PAS), (C) tuft cells (DCAMKL1, red) and (D) paneth cells (lysozyme, green) and epithelial cells (E-cadherin, blue). For (D) tissues were collected 2 hours post-EdU injection (200μg/mouse) and also stained for EdU+ cells (red). The number of EDU+ cells/crypt was quantified in (E). Images show representative villi from each genotype, bar graphs show quantitation of the indicated cell type. Each data point represents an average of >30 villi sampled per mouse (n=4). Error bars represent SD between samples. For panels A-E statistical analysis was performed using a student t-test, ns = non-significant. (F) Intestinal contractility was determined for longitudinally orientated intestinal sections using an organ bath. Following equilibration, the average cyclic height, contractility and tension of spontaneous contacts were then determined over 10 minutes. Statistical analysis was performed using Mann-Whitney test; p<0.01**, ns = non-significant, as data did not assume normal distribution. Related to Figure 3.

**Supplementary Figure 4.**
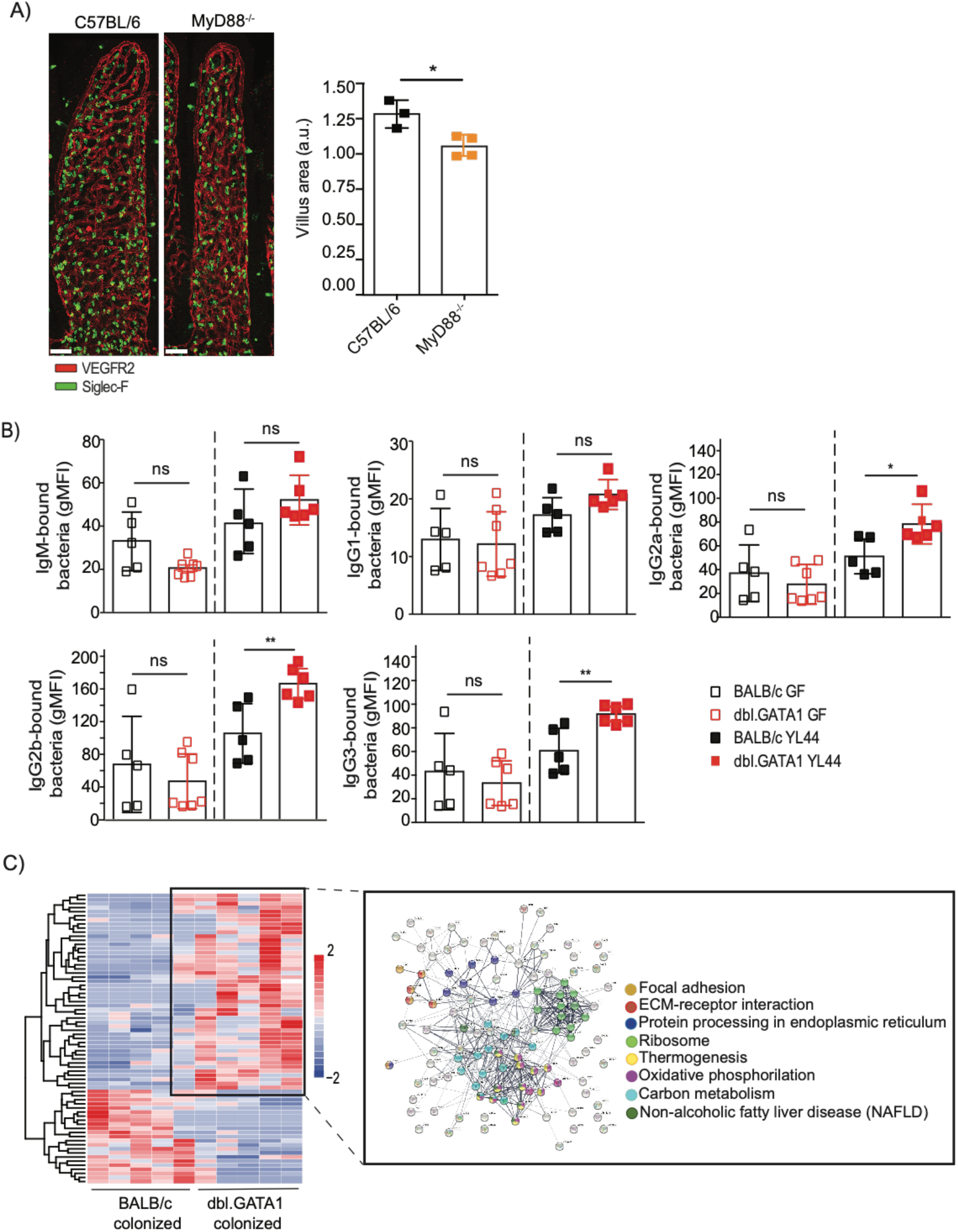
Eosinophil function is driven by microbiota. (A) Quantification of mean villous surface area from various SI regions calculated from the vascular cage of whole-mount tissues from C57BL/6 wild type mice vs. MyD88−/− mice SPF mice stained for eosinophils (Siglec-F, green) and blood endothelial cells (VEGFR2, red). Images show representative villi from a single mouse. Each bar graph represents data mean ± SD of n=3-4 mice with individual data points representing the mean surface area of at least >30 randomly sampled villi per mouse obtained from the proximal jejunum. Statistical analysis was performed using a student-test p<0.01** Germ-free (GF) BALB/c and Δdbl.GATA1 mice were monocolonized with *Akkermansia muciniphila* YL44 and serum collected 2 weeks later. The presence of *A. muciniphila* reactive serum immunoglobulin (Ig) was determined by flow cytometry. Data are representative of two independent experiments. Each bar graph represents data mean μ ± SD of n=5-7 mice with individual data points representing a single mouse. Dashed lines show GF and monocolonized mice separation for statistical analysis which was performed using a student t-test p<0.05*, p<0.01**, N.S. non-significant. (C) Heat map of proteins differentially expressed (log2FD > 1, p<0.05) in the jejunum fraction of Δdbl.GATA1 vs. BALB/c mice. STRING-db (KEGG pathways) analysis shows protein-protein interaction pathways enriched in the Δdbl.GATA1 mice. Related to Figure 4.

